# A Stomata Classification and Detection System in Microscope Images of Maize Cultivars

**DOI:** 10.1101/538165

**Authors:** Alexandre Hild Aono, James Shiniti Nagai, Gabriella da Silva Mendonça Dickel, Rafaela Cabral Marinho, Paulo Eugênio Alves Macedo de Oliveira, João Paulo Papa, Fabio Augusto Faria

## Abstract

Research on stomata, i.e., morphological structures of plants, has increased in popularity in the last years. These structures (pores) are in charge of the interaction between the internal plant system and the environment, working on different processes such as photosynthesis and transpiration stream. Besides, a better understanding of the pore mechanism plays a significant role when exploring the evolution process, as well as the behavior of plants. Although the study of stomata in dicots species of plants has advanced considerably in the past years, there is little information about stomata of cereal grasses. Also, automated detection of these structures have been considered in the literature, but some gaps are still uncovered. This fact is motivated by high morphological variation of stomata and the presence of noise from the image acquisition step. In this work, we propose a new methodology for automatic stomata classification and a new detection system in microscope images for maize cultivars. We have achieved an approximated accuracy of 97.1% in the identification of stomata regions using classifiers based on deep learning features, which figures out as a nearly perfect classification system.

## 1. Introduction

Stomata have probably received more attention than any other single vegetative structure in plants [1]. Regulating gas exchange between the plant and the environment[2], these structures stand for small pores on the surfaces of leaves, stems and parts of angiosperm flowers and fruits [3, 4], and they are formed by a pair of specialized epidermal cells (guarder cells) that are located in the surface of aerial parts of most higher plants [1]. Due to the controlling of the exchange of water vapor and CO^2^ between the interior of the leaf and the atmosphere [3], the photosynthesis, transpiration stream, nutrition and the metabolism of land plants are in different ways related to the opening and closing movements of the stomata [4, 1]. Furthermore, Hetherington and Woodward [3] pointed out that the acquisition of stomata and an impervious leaf cuticle are considered the key elements in the evolution of advanced terrestrial plants, allowing the plant to inhabit a range of different, often fluctuating, environments but still control water content.

The stomatal movements distinguish such structures from other pores found in plant organs, e.g., pneumathodes, hydathodes, lenticels, and the breathing pores found in the thalli of liverworts [1]. The control of stomatal aperture requires the coordinated control of multiple cellular processes [3] and its morphogenesis is affected by several environmental stimuli, such as relative humidity, temperature, the concentration of atmospheric carbon dioxide, light intensity, and endogenous plant hormones [2, 3, 1]. Global warming, for example, could increase leaf transpiration and soil evaporation, and as consequence leaf stomata movements can control plant water loss and carbon gain under this water stress condition [5]. Stomatal aperture might also represent an initial response to both plants and human pathogenic bacteria [2]. In plants, it has been reported that microscopic surface openings serve as passive ports of bacterial entry during infection and the stomatal closure is part of a plant innate immune response to restrict bacterial invasion [6].

The number of pores per unit area varies not only between species but also within species because of the influence of environmental factors during growth, leaf morphology and genetic composition [4, 1]. In general, it happens due to the influence on cell size [4], e.g., smaller guarder cells are usually associated with higher stomatal frequencies [1]. Besides stomata differentiation is a process that occurs together with the development of plant organs and, therefore, counts of stomata per unit area carried out at different stages in leaf development will differ [4]. Another characteristic with great variation concerns the spacing of stomata, which may be relatively evenly spaced throughout a leaf, located in regular rows along the length of a leaf, or they may be clustered in patches [1].

Since the types of stomatal configuration are profoundly different, the study and identification of these pores are vital points to understand several mechanisms of plants. Haworth et al. [7] also stated that it might be reasonable to assume that stomatal structures have played a significant role in plant evolution over the last 400 million years. Nevertheless, the examination of stomata from microscope images involves manual measurement and is highly dependent on biologists with expert knowledge to correctly identify and measure stomatal morphology [8].

Even with the apparent relevance of these structures, a recent study [9]indicated that, surprisingly, we still know little about stomata of cereal grasses. These grasses are extremely important since they provide the majority of calories consumed by humans either directly through the consumption of grains or indirectly through animals fed a diet of grains and forage[10]. Hepworth et al. [9] highlighted that the stomatal complexes in grasses differ of the dicots in many ways, e.g., the guard cells of dicots are kidney-shaped and form stomata that are scattered throughout the epidermis in a less orderly pattern, while stomatal configuration of grasses develop in parallel rows within defined and specific epidermal cell files [9].

In this scenario, in order to assist the biological community to perform stomata studies, we proposed an automated strategy for stomata detection and classification in microscope images using machine learning techniques. Our work is seminal in a sense it is less time-consuming when examining stomatal behavior, thus enabling biologists to use more information from the images and study a broader range of stomata. In this work, we employed microscope images of maize, which represent the most produced and consumed cultivars in the world. As far as we are concerned, we have not observed any similar work concerning maize cultivars.

The remainder of this work is organized as follows. Section 2 and 3 present the related works and the proposed approach, respectively. Section 4 discusses the methodology, while Section 4 presents the experiments. Finally, Section 6 states conclusions and future works.

## 2. Related Works

The research of stomata image processing started in the 80’s. Recognized as possible pioneers, Omasa and Onoe [11] proposed a technique for measuring stomata characteristics in grayscale images using Fourier Transform and threshold filters for image processing and segmenting [8]. More recently, Sanyal et al. [12] compared tomato cultivars using several morphological characteristics, including stomata measures. Microscope images of different varieties were obtained using a scanning electron microscope, and the segmentation was performed using a watershed algorithm resulting in one stomata per image, followed by morphological operations (e.g., erosion and dilation) and Sobel kernel filters to remove noise and obtain stomatal boundaries. Using 100 images of tomato cultivars and a multilayer perceptron algorithm, the proposed approach achieved 96.6% of accuracy.

Jian et al. [13] aimed at estimating stomata density using three different regions of *Populus Euphratica* leaves. For image processing purposes, an object-oriented classification method was used with parameters such as scale, compactness, and shape. Such an approach presented high accuracy when compared to human-based count, showing advantages over the traditional method to extract the stoma information. Aiming the constant growth and development of stomata image processing studies, Higaki et al. [14] published the “Live Images of Plant Stomata LIPS” database. In other work, Higaki and Takumi [15] presented a semi-automatic stomata region detection approach using ImageJ software [16] and a Clustering-Aided Rapid Training Agent-based algorithm [17].

Oliveira et al. [18] proposed an approach solely based on morphological operations. Initially, a Gaussian low-pass filter was employed to preprocess the images and remove noise. Further, reconstruction operations (e.g., opening and closing) were applied to highlight stomata regions, which were counted based on background intensity differences. As a result, the work reported recognition rates of around 94.3%.

Laga et al. [19] introduced a supervised method for stomata detection based on morphological and structural features. To fulfill such purpose, 24 microscope images were obtained and filtered by normalization together with a Gaussian filter. The images were manually segmented and the width and height parameters extracted. The authors reported results close to a manual counting approach. Later on, a patent for stomata measurement using Gaussian filtering and morphological operations was registered by Awwad et al. [20].

Duarte et. al.[21] proposed a method to count stomata in microscope images automatically. Initially, the images were converted from RGB to CieLAB to select the best channel for analysis. Wavelet Spot Detection and morphological operations performed the stomata detection step, with results nearly to 90.6% of recognition accuracy.

Jayakody et al. [8] proposed an automated stomata detection and pore measurement system for grapevines. The approach employed a Cascade Object Detection (COD) algorithm with two main steps: (i) first, the COD classifiers are trained using stoma and non-stoma images, and then (ii) a sliding window over the microscope images was used to identify stomata inside it. After its detection, the pore measurement step was performed using binary segmentation and skeletonization with ellipse fitting, for further estimating pore measurements. The authors reported 91.6% of recognition rate.

## 3. Proposed System

This section introduces the approach for stomata classification and detection employed in the work.

### 3.1. Overview

The proposed approach is composed of two different process: (i) stomata detection and further (ii) classification. Figure 1 depicts an overview of the proposed approach.

**Figure 1:**
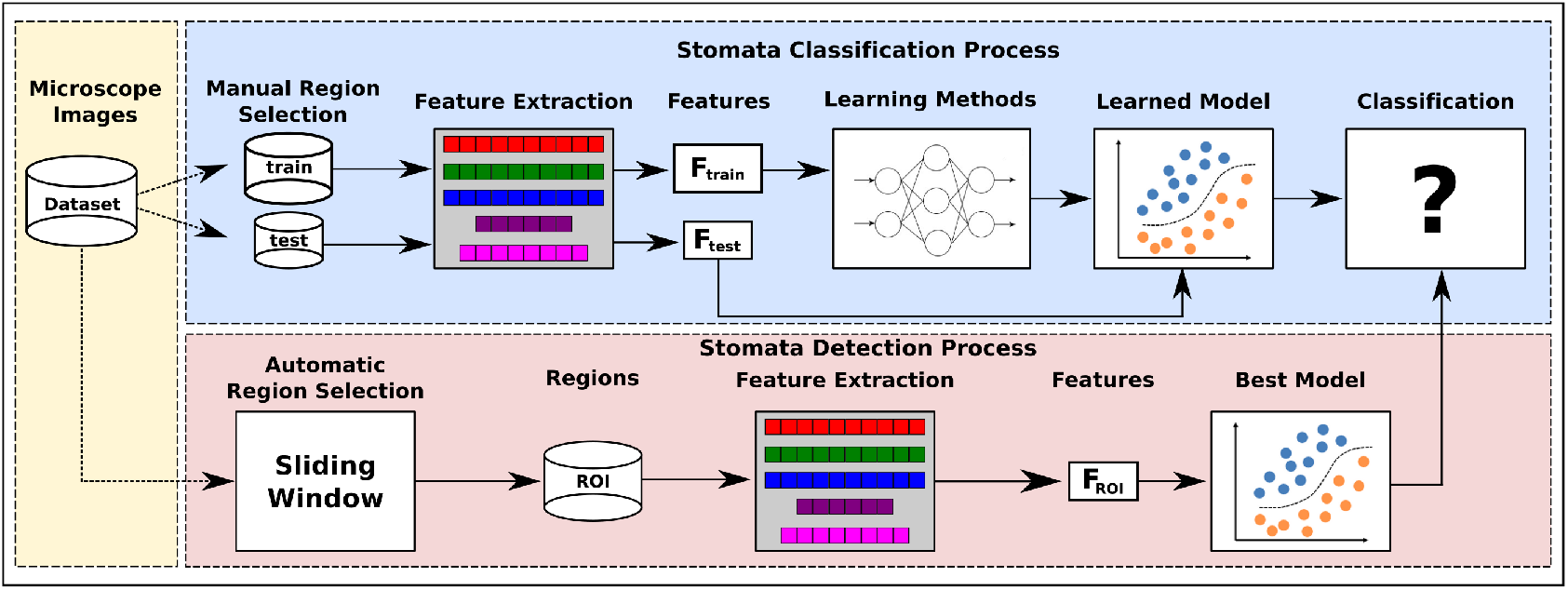
An overview of the proposed stomata classification and detection system.

In the stomata classification process, the first step is to manually collect and label a subset of stomata and non-stomata regions from the microscope images dataset, creating two disjoint sets of subimages, i.e., *train* and *test*. Such sets are subjected to an image descriptor that encodes the visual properties of the subimages into feature vectors (i.e., *F*_*train*_ and *F*_*test*_ for the *train* and *test* sets, respectively). Further, the feature vectors *F*_*train*_ are used as input for a learning method, thus creating a learned model for stomata classification purposes. Finally, each feature vector *F*_*test*_ is then classified by this learned model. In the classification process, different image descriptors and learning methods are evaluated through a *k*-fold cross-validation protocol, and the best model is adopted to detect stomata regions on the next step.

Regarding the stomata detection process, a sliding window is used on each microscope image from the entire dataset to create a set of regions of interest (*ROI*), which are subjected to an image descriptor resulting in the feature vectors (*F*_*ROI*_). Finally, each *F*_*ROI*_ is classified by the best model, i.e., a tuple (learning method + image descriptor) computed in the classification process.

### 3.2. Stomata Classification Process

The first step for identifying stomata structures is the manual selection of a set of subimages containing stomata or other plant structures, labeled as non-stomata. Due to the differences between stomata size in distinct microscope images, we adopted a region/window of dimension 151 × 258 pixels. We observed that such size is enough to include all stomata regions from the dataset images. Therefore, a total of 1, 000 subimages of each class (i.e., stomata and non-stomata) were selected to compose the new dataset.

Once the dataset has been created, the next step is to extract visual properties from the subimages using image descriptors. In this work, we evaluated eleven different image descriptors, as described below:

#### 3.2.1. DAISY

the descriptor relies on gradient orientation histograms. For an input image, orientation maps are calculated based on quantized directions using Gaussian kernels. The final descriptor concerns the values from these convolved maps located on concentric circles centered on a location. The amount of Gaussian smoothing is proportional to the radius of the circles [22].

#### 3.2.2. Histogram of Oriented Gradients (HOG)

feature descriptor based on the creation of histograms with gradient orientation using their magnitude in specific portions of an image [23]. The local shape information is described by the distribution of gradients in different orientations [24].

#### 3.2.3. GIST

the descriptor focuses on the shape of the scene itself, i.e., on the relationship between the outlines of the surfaces and their properties, ignoring the local objects in the scene and their relationships [25]. The approach does not require any form of segmentation and is based on a set of perceptual dimensions (naturalness, openness, roughness, expansion, ruggedness) [24].

#### 3.2.4. Haralick Texture Features

at first, a gray-level co-occurrence matrix is computed considering the relation of each voxel with its neighborhood. Using different statistical measures (e.g., entropy, energy, variance, and correlation), texture properties are encoded from the image into feature vectors [26].

#### 3.2.5. Local Binary Patterns (LBP)

it computes a local representation of texture based on the comparison of each pixel with its neighborhood. A threshold for such comparison is defined and an output image is produced with the binary to decimal values conversion. Further, a histogram is created as the final descriptor [27].

#### 3.2.6. Deep Convolutional Neural Network

a typical convolutional network is a fully-connected network where each hidden activation is computed by multiplying the entire input by weights in a given layer [28]. In this technique, a connection between traditional optimization-based schemes and a neural network architecture is considered, where a separable structure is introduced as a reliable support for robust deconvolution against artifacts [29]. Once we do not have available a large scale of images to train a deep learning architecture from scratch, a good alternative is to use the transfer learning [30]. Usually, the networks are pre-trained over ImageNet dataset [31], for further adding other layers according to the target application. The last layer can be used for feature extraction purposes (image descriptor). In this work, we adopted six different architectures: (i) DenseNet121 [32], (ii) InceptionResNetV2 [33], (iii) InceptionV3 [34], (iv) ModbileNet [35], (v) NasNet [36], and (vi) VGG16 [37].

Concerning the machine learning techniques, we used three different approaches: (i) Support Vector Machine [38] (SVM), (ii) Multilayer Perceptron [39] (MLP), and (iii) Adaboost [40]. The best tuple (i.e., learning method + image descriptor) will be then employed to label the new stomata regions on the next process. Figure 2 shows the steps of the stomata classification process proposed in this work.

**Figure 2:**
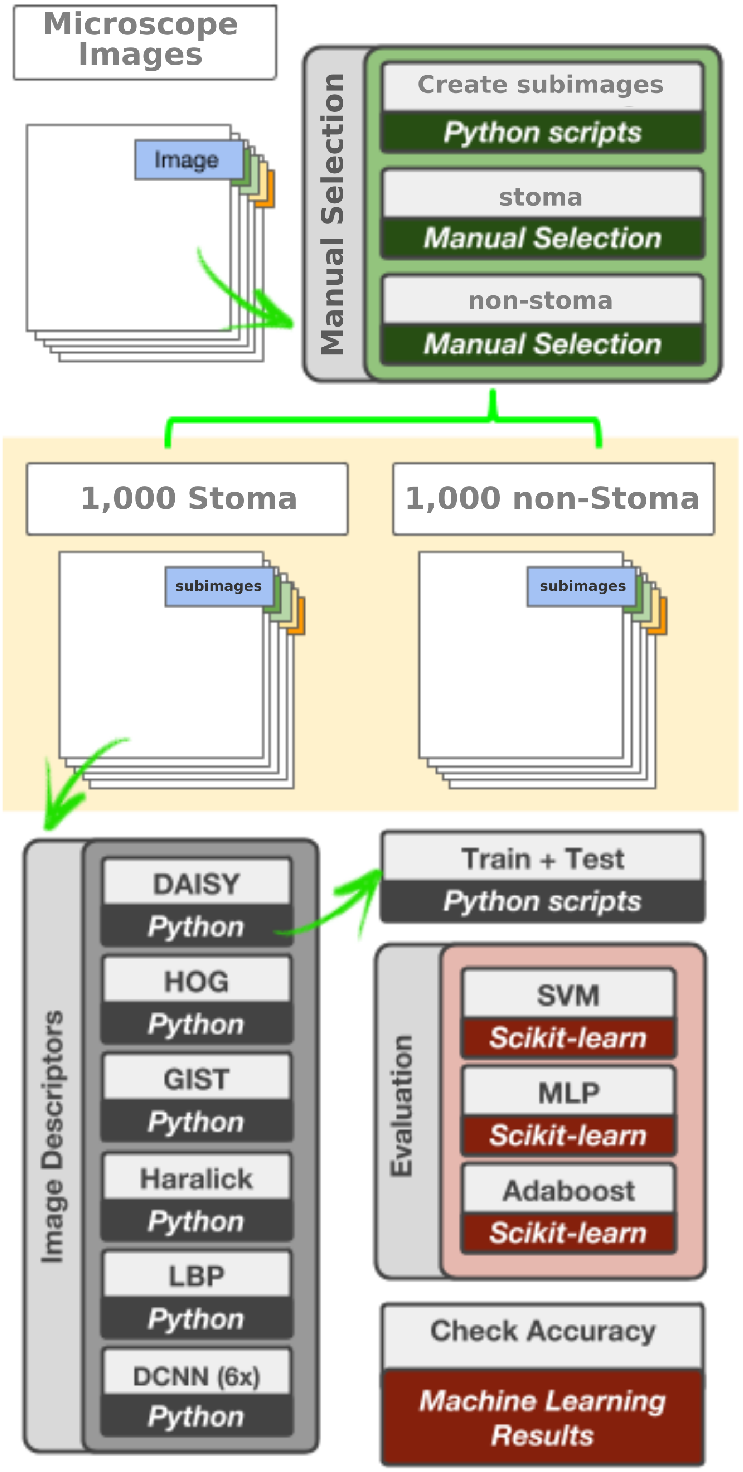
In-depth explanation of the stomata classification process.

### 3.3. Stomata Detection Process

Figure 4 depicts the methodology for stomata detection, which is divided into the following steps:

#### 3.3.1. Dataset

A dataset with stoma and non-stoma subimages (Figure 3) was created through a manual selection task from microscope images.

**Figure 3:**
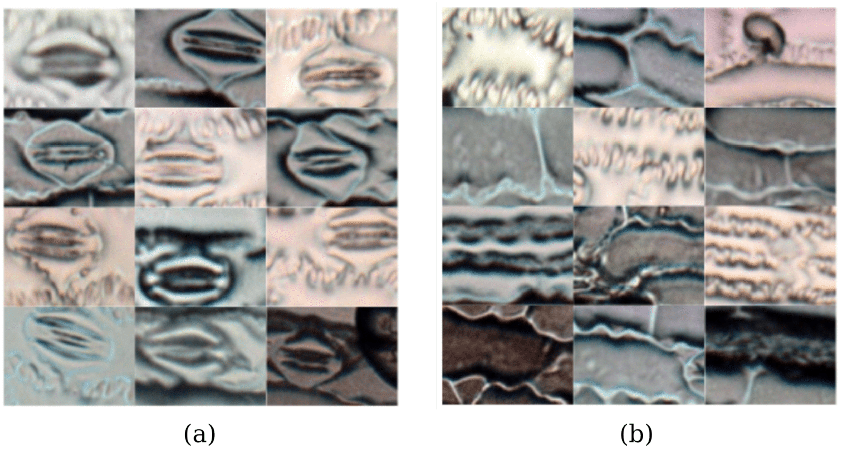
Examples of: (a) stoma and (b) non-stoma subimages/regions, which were manually selected and labeled in this work.

**Figure 4:**
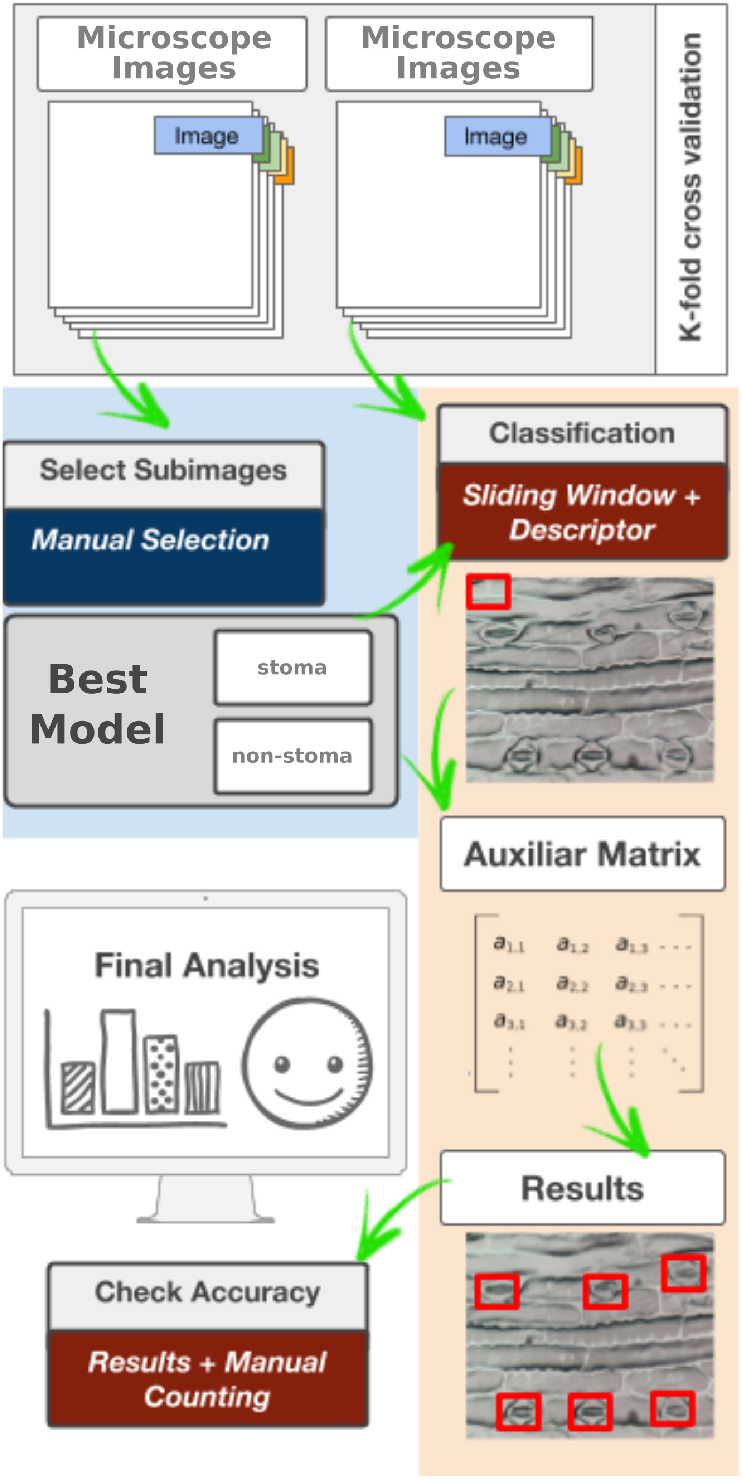
In-depth explanation of the stomata identification process.

#### 3.3.2. Feature extraction

Once the best descriptor has been found on the stomata classification process, the features of the new dataset are generated and stored into a table with the labels of each category (stoma or non-stoma).

#### 3.3.3. Creation of the learned model

The descriptors were evaluated using three different learning methods: SVM, MLP, and Adaboost. Based on the best effective results achieved by each learned model (i.e., a tuple composed of a aescriptor + the learning method), the most appropriate learned model is then selected to label the subimage in next step.

#### 3.3.4. Sliding window iteration

Using a window of 151 × 258 pixels, an iteration over the microscope images is performed, and for each generated subimage, a label (stoma or non-stoma) is obtained using the best-learned model. Due to the possible separation of stoma structures, the windows were created with a stride of 100 pixels in both columns and rows.

#### 3.3.5. Selection of positive regions

Based on the previous classification, an auxiliary matrix is filled in order to enable the posterior identification of stoma regions. Pixels with a positive occurrence for stoma are separated from the rest of the image, for the further analyzes of such regions.

## 4. Experimental Setup

This section describes the dataset design, the technologies, and evaluation protocol used in this work.

### 4.1. Image Dataset

Regarding optical microscope investigation, it has been necessary to separate the epidermis from the remainder of the leaf itself to get a clear view of the cell walls and the shape of the stomata [41]. Herein cyanoacrylate glue was applied to the microscope slide in order to obtain an impression of the sheet surface to be captured using a camera attached to a microscope. Leaves were sampled from 20 *Zea mays* cultivars (maize) granted by Nidera Sementes company (Uberlândia-MG), producing a total of 200 microscope images with different dimensions such as 2, 565 × 3, 583, 2, 675 × 3, 737, and 2, 748 × 3, 840.

The selected species then were treated with colchicine [42] to change their ploidy and cell morphology for further studies. Due to the plant ploidy specificity, different images might have different stomata sizes and width. Besides, as previously mentioned, stomata differentiation is a process that occurs together with the development of plant organs, and herein plants with different ages were used, and a clear distinction of the images and plant morphologies can be visualized in Figure 5. In these microscope images, different types of noise and artifacts can be observed as well, as depicted in Figure 6, thus highlighting the challenges faced in this work.

**Figure 5:**
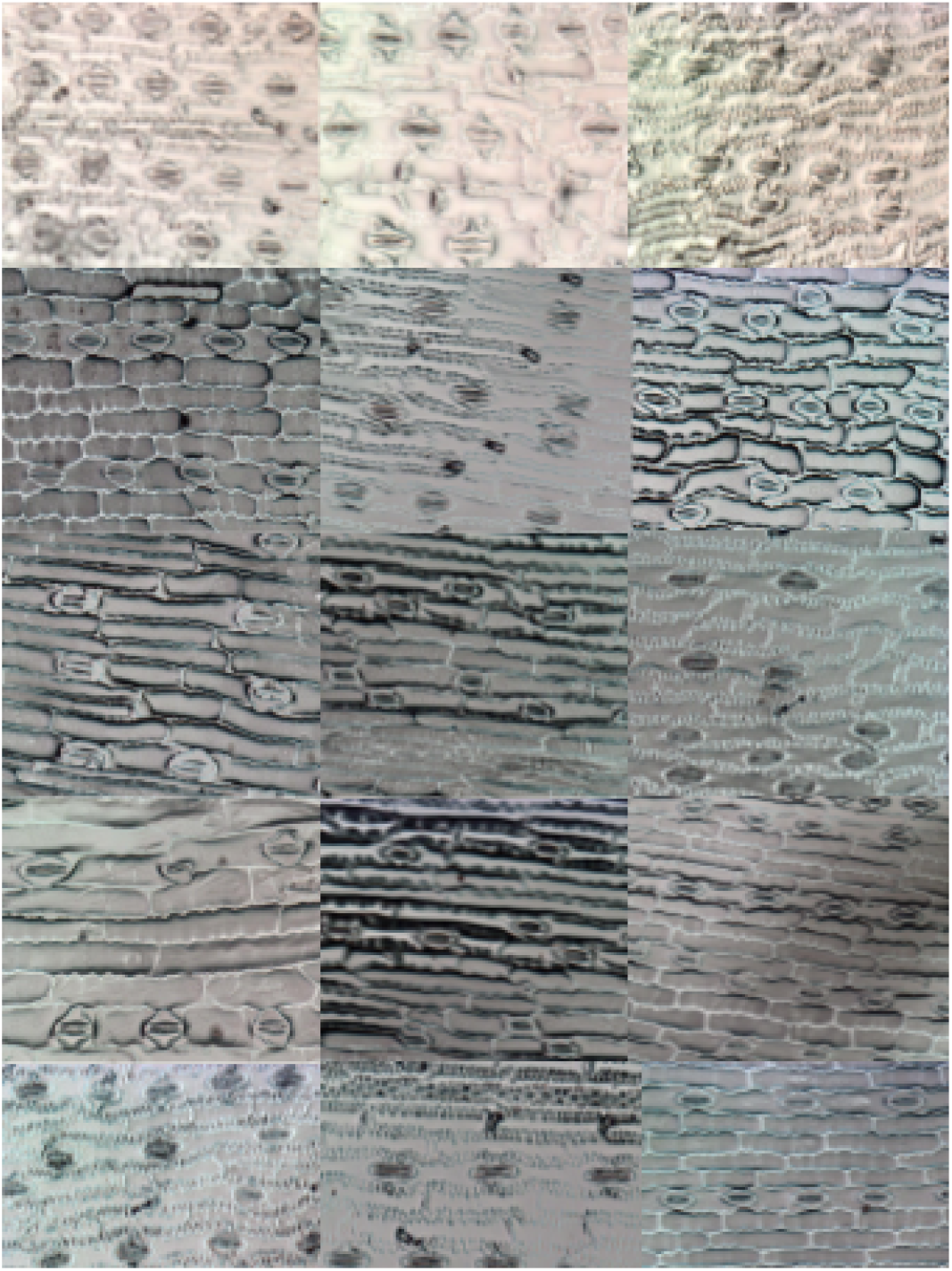
Samples from the microscope images of Maize Cultivars used in this work.

**Figure 6:**
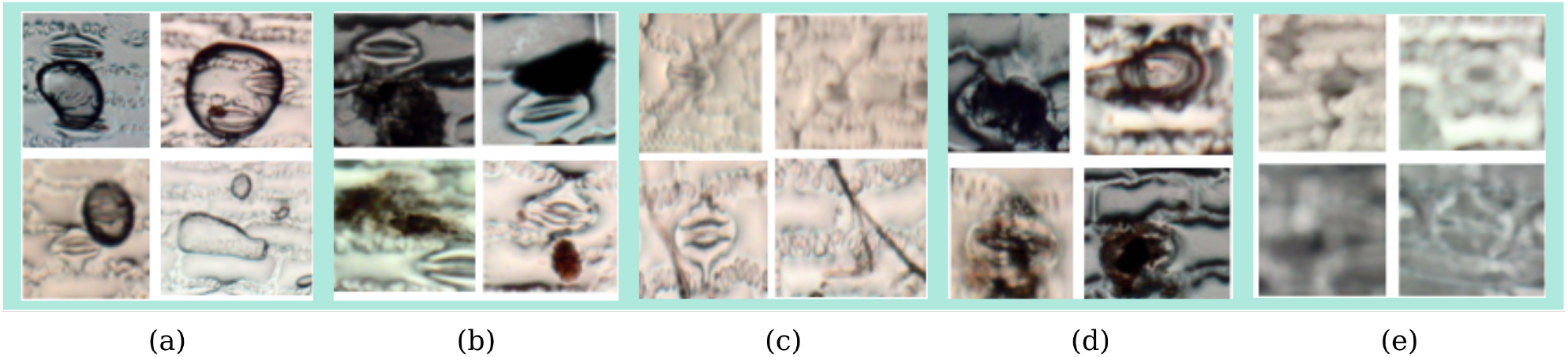
Different types of noise present in the microscopic images: (a) the usage of cyanoacrylate glue can generate air bubbles; (b) the microscope might capture leaves residuals; (c) the leaves might bend and create grooves in the image; (d) degraded stomata due to biological factors; and (e) low image quality due to equipment limitations.

In the experiments, the dataset with 200 microscope images was submitted to the 5-fold cross-validation protocol, i.e., four parts of the dataset compose the training set (160 images), and one part belongs to the test set (40 images). This process is repeated five times. Therefore, in the *stomata classification task*, for each microscope image, 5 stoma and 5 non-stoma regions/sub-images have been manually select to compose training and test sets in an overall of 2, 000 sub-images.

Concerning the *stomata detection task*, respecting the separation of the disjoint sets of the 5-fold cross-validation protocol, each training set created in the stomata classification task is maintained with 1, 600 sub-images. However, the test sets are generated by a sliding window operation. Hence, for each one of the 40 test images, between 876 and 963 regions/sub-images were selected by a sliding window iteration, resulting in approximately 44, 000 sub-images per test set, in an overall of 217, 866 sub-images for the five runs.

### 4.2. Programming Environment and Libraries

All approaches considered in this paper were executed on a personal computer with 2.7GHz Intel Core i7-7500U 2.7GHz Intel Core i7-7500U with 16GB of RAM and NVIDIA GeForce 940MX 4GB graphic card. Similarly, the programming language used in this work was Python2 with the following libraries: scikit-learn [43], pyleargist, scikit-image[44], opencv [45], keras[46] and tensorflow[47]. A considerable part of the libraries was mostly used for feature extraction and deep learning methods purposes.

### 4.3. Evaluation Protocol

To assess the accuracy of the proposed approach for classifying and identifying stomata regions, we employed a *k*-fold cross-validation with *k* = 5. The classified images represent the test set and the sub-images used to create the learned model were extracted from the training set. A manual count was also performed for each image to evaluate the final results using all windows generated, including the overlapped regions.

## 5. Results and Discussion

This section discusses the experiments performed to validate the proposed approach.

### 5.1. Stomata Classification Task

In this first experiment, we performed a comparative analysis among five image descriptors (HOG, GIST, DAISY, LBP, and Haralick) and three learning methods (Adaboost, MLP, and SVM) for the stomata classification task. The effectiveness is measured in terms of the mean accuracy considering the 5-fold cross-validation protocol.

As one can observe in Table 1, the best results were achieved by descriptors purely based on gradient information (HOG and DAISY)^1^. HOG descriptor with MLP (HOG+MLP) and DAISY descriptor with Adaboost (DAISY+Adaboost) achieved 96.0% of mean accuracy. In a comprehensive comparison among all image descriptors, HOG descriptor was the most effective with a mean accuracy of 94.7%, which can be justified by the specific shape of the stoma when compared to other parts. Therefore, this fact can show us that shape is perhaps the most essential visual property for the target application. Although GIST is a shape descriptor, its way of dealing with visual properties globally (holistic) may explain its poor performance in such images.

**Table 1:** Mean Accuracy of the classifiers based on image descriptor features for the stomata classification task.

| Learning Method | HOG | GIST | DAISY | LBP | Haralick |
| --- | --- | --- | --- | --- | --- |
| Adaboost | 93.0 | 79.0 | **96.0** | 88.0 | 87.0 |
| MLP | **96.0** | 81.0 | 92.0 | 85.0 | 80.0 |
| Linear SVM | **95.0** | 81.0 | 80.0 | 89.0 | 86.0 |
| **Mean** | 94.7* | 80.3 | 89.3 | 87.3 | 84.3 |

Since deep learning techniques are on the spotlight due to their outstanding results in a number of applications, we also considered them in this work. Table 2 presents the effectiveness results of six different deep learning architectures (DenseNet121 – DenseNet, Inception-ResNetV2 – IResNet, InceptionV3 – Inception, MobileNet, NasNet, and VGG16) using three learning methods (Adaboost, MLP, and SVM) concerning the stomata classification task.

**Table 2:** Mean Accuracy of the experiments based on deep learning features for the stomata classification task.

| Classifier | DenseNet | IResNet | Inception | MobileNet | NasNet | VGG16 |
| --- | --- | --- | --- | --- | --- | --- |
| AdaBoost | 95.0 | 96.0 | 90.0 | 96.0 | 91.0 | **99.0** |
| MLP | 98.0 | 94.0 | 88.0 | 98.0 | 95.0 | **100.0** |
| Linear SVM | 80.0 | 94.0 | 91.0 | 98.0 | 95.0 | **100.0** |
| **Epochs** | 10 | 13 | 6 | 7 | 16 | 6 |
| **Mean** | 91.0 | 94.7 | 89.7 | 97.3 | 93.7 | 99.7* |

As one can observe, information based on deep learning features outperformed the handcrafted ones (Table 1), except for HOG descriptor. In this experiment, the classifiers using VGG16 features achieved the best results with 100% of mean accuracy for almost all three learning techniques considered in this work for the stomata classification task.

### 5.2. Stomata Detection Task

In this experiment, the classifier based on VGG16 features with Support Vector Machines (SVM+VGG16) was adopted for the stomata detection task since it obtained the best results in the stomata classification task. Using the sliding window approach to generate possible stomata regions, we have created between 876 and 963 regions/sub-images for each microscope image (overall of 217, 866 sub-images) for the further application of a 5-fold cross-validation protocol.

Table 3 summarizes the effectiveness results considering the classifier SVM+VGG16. The number of detected stoma regions are compatible with the manual counting, which shows a good performance of the proposed approach. Besides, all folds presented similar effectiveness with around 97.1% of detected stoma regions, i.e., 11, 388 stomata out of the 11, 734 ones present in the dataset. It is also important to clarify that the results achieved in this paper are better than the ones recently reported by Jayakody et al. [8], which obtained an overall accuracy of 91.6% of detected regions.

**Table 3:** Effectiveness results of the classifier (SVM+VGG16) for sliding window classification.

| Fold | # Stoma Manual Counting | # Detected Stoma Regions | Total of Regions | # True Positives | # False Positives |
| --- | --- | --- | --- | --- | --- |
| 1 | 2,244 | 2,189 (97.5%) | 43,524 | 5,094 | 107 (0.02%) |
| 2 | 2,374 | 2,300 (96.9%) | 43,458 | 5,307 | 159 (0.03%) |
| 3 | 2,428 | 2,316 (95.4%) | 43,524 | 5,506 | 153 (0.03%) |
| 4 | 2,279 | 2,213 (97.1%) | 43,680 | 5,596 | 60 (0.01%) |
| 5 | 2,409 | 2,370 (98.4%) | 43,680 | 5,463 | 49 (0.01%) |
| Mean | — | 2277.6 (97.1%) | — | 5,393.2 | 105.6 (0.02%) |
| Overall | 11,734 | 11388 | 217866 | — | — |

Once the stomata region candidates have been detected in a microscope image (Figure 7(a)), an auxiliary matrix was created to encode the stomata region occurrence (Figure 7(b)), and then a merging between microscope image and auxiliary matrix was performed (Figure 7(c)). Finally, all stomata are identified in the microscope image, as depicted in Figure 7(d).

**Figure 7:**
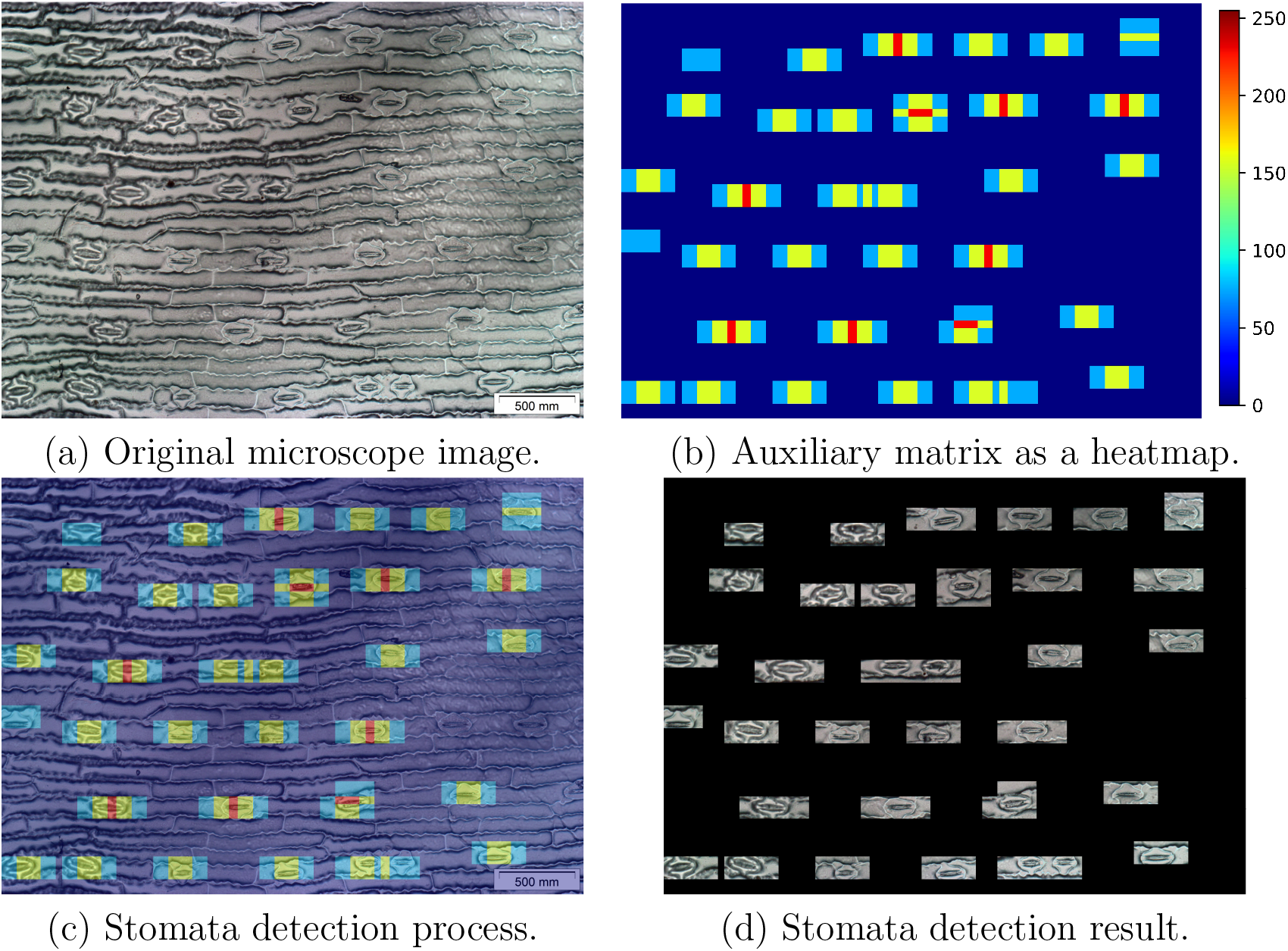
Pos-processing of a microscope image.

We have also analyzed the quality of the effectiveness results. Figure 8 shows the hit and miss-classification results achieved by the proposed system. It is essential to observe that regions/sub-images with low quality have also been correctly classified as containing stoma, as depicted in Figure 8(a). This fact corroborates the usage of the VGG16 features for the stomata detection task. Miss classified regions can be visualized in Figure 8(b). Most of these regions/sub-images represent plant structures that are similar to stomata.

**Figure 8:**
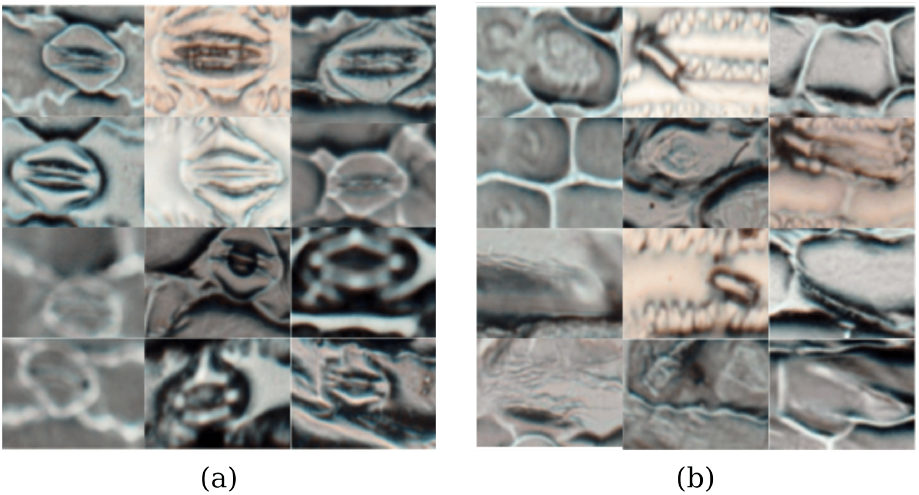
Examples of the stomata classification results: (a) true positive sub-images and (b) false positive sub-images.

## 6. Conclusions

Leaves microscope images contain relevant information about plant morphology and can be used for studying specific characteristics of metabolic pathways and different biological processes. A vegetative structure that has received considerable attention concerns the so-called stoma (in the plural, stomata), which stands for small pores on the surfaces of aerial parts of most higher plants (e.g., leaves, stems and pieces of angiosperm flowers and fruits). Stomata are responsible by many functionalities such as (i) exchange of water vapor and CO^2^ between the interior of the leaf and the atmosphere; (ii) photosynthesis; (iii) transpiration stream; (iv) nutrition; and (v) metabolism of land plants. Therefore, the understanding of the stomata is of great importance in the exploration of the evolution and behavior of plants.

In this work, we proposed a stomata classification and identification approach in microscope images of maize cultivars. Herein we have evaluated different extraction techniques (image descriptor and deep learning) and learning methods (Adaboost, MLP, and SVM) concerning the task of correctly classifying stomata regions. In the experiments, the proposed approach achieved a mean accuracy of 96% using HOG+MLP, and a mean accuracy of 100% with VGG16 features using Support Vector Machines (VGG16+SVM).

Regarding the stomata detection task with a sliding window approach for generating all possible regions/sub-images from the microscope images, the proposed approach detected 97.1% of the stomata regions in 200 microscopes. This fact could show us that the proposed approach using deep learning features might be an appropriate solution for the target application.

As future work, we intend to develop a computational toolkit to support the specialists in the biology area in their research.

## Acknowledgements

The authors thanks the support of the Brazilian scientific funding agency CNPq through projects #408919/2016-7 and #307066/2017-7, São Paulo Research Foundation FAPESP grants #2016/19403-6, #2014/12236-1, and #2013/07375-0, as well as the support of NVIDIA Corporation for the donation of the GPUs used for this research.

## Footnotes

1 The values in bold stand for the best descriptor per classifier. Symbol ‘⋆’ denotes the best overall result.

